# Effects of tDCS on motor learning and memory formation: a consensus and critical position paper

**DOI:** 10.1101/064204

**Authors:** Ethan R Buch, Emiliano Santarnecchi, Andrea Antal, Jan Born, Pablo A Celnik, Joseph Classen, Christian Gerloff, Mark Hallett, Friedhelm C Hummel, Michael A Nitsche, Alvaro Pascual-Leone, Walter J Paulus, Janine Reis, Edwin M Robertson, John C Rothwell, Marco Sandrini, Heidi M Schambra, Eric M Wassermann, Ulf Ziemann, Leonardo G Cohen

## Abstract

Motor skills are required for activities of daily living. Transcranial direct current stimulation (tDCS) applied in association with motor skill learning has been investigated as a tool for enhancing training effects in health and disease. Here, we review the published literature investigating whether tDCS can facilitate the acquisition and retention of motor skills and adaptation. A majority of reports focused on the application of tDCS with the anode placed over the primary motor cortex (M1) during motor skill acquisition, while some evaluated tDCS applied over the cerebellum during adaptation of existing motor skills. Work in multiple laboratories is under way to develop a mechanistic understanding of tDCS effects on different forms of learning and to optimize stimulation protocols. Efforts are required to improve reproducibility and standardization. Overall, reproducibility remains to be fully tested, effect sizes with present techniques are moderate (up to d= 0.5) (Hashemirad, Zoghi, Fitzgerald, & Jaberzadeh, 2016) and the basis of inter-individual variability in tDCS effects is incompletely understood. It is recommended that future studies explicitly state in the Methods the exploratory (hypothesis-generating) or hypothesis-driven (confirmatory) nature of the experimental designs. General research practices could be improved with prospective pre-registration of hypothesis-based investigations, more emphasis on the detailed description of methods (including all pertinent details to enable future modeling of induced current and experimental replication) and use of post-publication open data repositories. A checklist is proposed for reporting tDCS investigations in a way that can improve efforts to assess reproducibility.

## 1 Introduction

Noninvasive brain stimulation (NIBS), most commonly repetitive transcranial magnetic stimulation (rTMS) and transcranial direct current stimulation (tDCS), have been used to modulate motor and cognitive functions in human subjects (Brunoni et al., 2012; Bütefisch, Khurana, Kopylev, & Cohen, 2004; Duque et al., 2007; Gangitano et al., 2002; Jahanshahi & Rothwell, 2000; Maeda, Keenan, Tormos, Topka, & Pascual-Leone, 2000; Marshall, Helgadottir, Molle, & Born, 2006; Pascual-Leone, Valls-Sole, Wassermann, & Hallett, 1994; Perceval, Floel, & Meinzer, 2016; Wassermann, Tormos, & Pascual-Leone, 1998) (Figure 1)^a^. It has been argued that rTMS and tDCS can either enhance or decrease excitability in targeted cortical regions depending on the parameters of stimulation employed (Chen et al., 1997; Galea, Jayaram, Ajagbe, & Celnik, 2009; Labruna et al., 2016; Woods et al., 2016) and the underlying intrinsic state of the stimulated brain networks (Dayan, Censor, Buch, Sandrini, & Cohen, 2013; Sandrini, Umilta, & Rusconi, 2011; Silvanto, Cattaneo, Battelli, & Pascual-Leone, 2008; Silvanto & Pascual-Leone, 2008).

**Figure 1.**
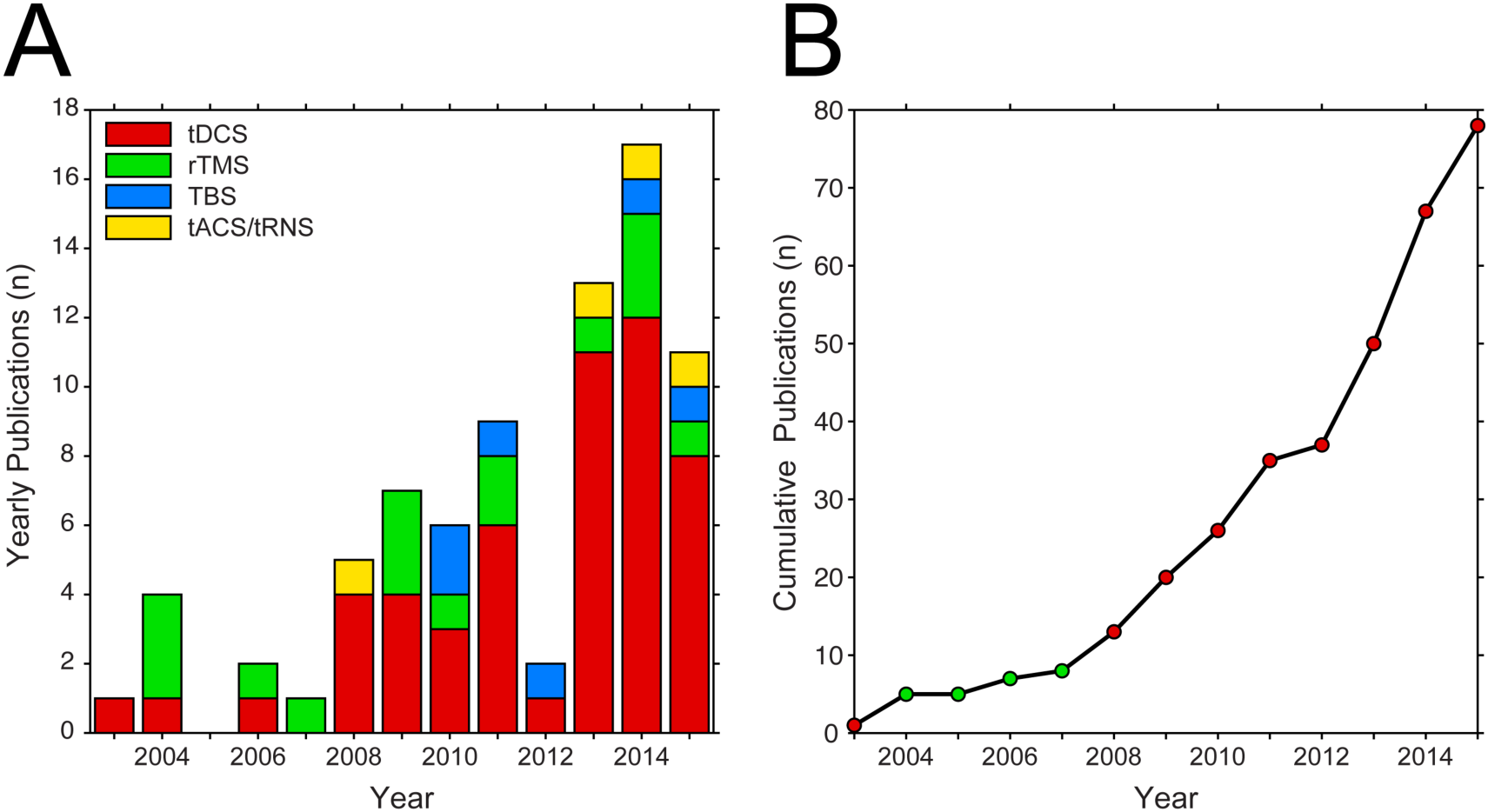
Publications of studies investigating NIBS-based enhancement of motor learning or memory formation. (A) Yearly publications grouped into categories of different noninvasive brain stimulation techniques. Categories consist of tDCS, rTMS (e.g. – 1, 5 or 10 Hz), TBS (e.g. - iTBS or cTBS), or tACS/tRNS. (B) Cumulative publications by year. Marker color indicates the majority stimulation type used in studies for that year.

tDCS has also been used as a tool to gain insight into brain-behavior interactions and to explore possible causal relationships between altered activity in relatively large regions of the brain and particular behaviors(Nitsche et al., 2008). More specifically, tDCS has been used to study effects on, and mechanisms of, motor learning (Antal et al., 2004; Galea, Vazquez, Pasricha, Orban De Xivry, & Celnik, 2011; Nitsche et al., 2003; Reis et al., 2009). In a previous consensus document, it was stated that “*Improved understanding of the involvement of a brain region in a type of behavior was followed by attempts to modify activity … to secondarily influence performance, learning and memory functions”* (Reis et al., 2008). Several recommendations from that paper have been advanced in the literature. For example, many studies have utilized multi-session rather than single-session tDCS application (Hashemirad et al., 2016), greater emphasis has been placed on monitoring long-term effects of tDCS on motor learning (Hashemirad et al., 2016), evidence of dissociation of tDCS effects applied to distinct brain regions on different stages of motor learning (Galea et al., 2011; Reis et al., 2015; Wymbs, Bastian, & Celnik, 2016) has begun to emerge, and mechanisms underlying tDCS effects are starting to be elucidated (Fritsch et al., 2010; H. I. Kuo et al., 2013; M. F. Kuo et al., 2008; Lang, Nitsche, Sommer, Tergau, & Paulus, 2003; Stagg, Bachtiar, & Johansen-Berg, 2011). Below, we summarize results from tDCS studies aiming to improve motor learning in healthy humans without performing a critical review of each individual investigation, discuss new challenges and limitations to be considered, and propose strategies to move forward.

### 1.1 Motor learning

The acquisition and retention of new motor skills, and adaptation of previously learned ones are fundamental to our daily lives (Debas et al., 2010). Commonly used skills such as typing or playing a musical instrument are acquired and improved through years of repetitive practice (Dayan & Cohen, 2011). This process remains adaptive throughout the lifespan, as the interaction between intrinsic (e.g. – body morphology, muscle strength, injury, etc.) and extrinsic (e.g. – tools, task constraints) factors require continuous updating of how we interact with an often changing environment (Wolpert, Diedrichsen, & Flanagan, 2011) undergoing consolidation (Muellbacher et al., 2002) and reconsolidation (Censor, Sagi, & Cohen, 2012). In the laboratory, motor learning is commonly explored using paradigms focusing on the acquisition and retention of new motor skills, or the adaptation of existing ones to environmental disruptions. Motor skill learning is typically achieved slowly with prolonged training, resulting in slow performance gains underpinned by an improved speed-accuracy relationship and/or a reduction in performance variability (Shmuelof, Krakauer, & Mazzoni, 2012). Conversely, motor adaptation is typically achieved over brief training periods, where performance levels are restored to prior maximums following exposure to an environmental perturbation (Shmuelof et al., 2012). Here, we focus on these most commonly studied types of motor learning, as both have been used as the substrate for neuromodulation. However, it should be kept in mind that the categorization of skill learning and adaptation is applied rather broadly and may engage error-dependent, use-dependent, reinforcement, and/or strategic learning to different extents (Krakauer & Mazzoni, 2011) with shared or independent underlying mechanisms.

From a behavioral standpoint, motor skill learning can be deconstructed into several component features that occur over different timespans. Learning is initiated by experience that is accrued over one or more practice or training periods (Dayan & Cohen, 2011). Performance improvements that occur over shorter time periods, such as within a single training session or day, are typically referred to as online learning (Reis et al., 2009). Over longer periods of time, such as over several hours, days or training sessions, motor memories may transition to a consolidation phase (Gais et al., 2007; Marshall & Born, 2007; Stickgold, 2005; Walker, Brakefield, Hobson, & Stickgold, 2003). Behavioral expressions of consolidation may include: (1) a greater resistance to interference caused by other learned skills (i.e. – stability)(Krakauer & Shadmehr, 2006); (2) observed performance improvements at re-test in the absence of additional practice (i.e. – offline gains) (Reis et al., 2009); or (3) reductions in performance decrements experienced with the passage of time (i.e. – retention) (Abe et al., 2011).

Even once acquired motor skills are consolidated and retained as stable, long-term motor memories they must maintain some capacity to be flexible and responsive to unpredictable biological or environmental changes that may occur in the future (Sandrini, Cohen, & Censor, 2015). Each time a given skill is executed, retrieval of these previously consolidated motor memories may initiate a cascade of plasticity mechanisms that enable their composition to be modified in order to maintain skill performance optimization over the long term (Censor, Buch, Nader, & Cohen, 2015; Censor, Dayan, & Cohen, 2014; Censor, Dimyan, & Cohen, 2010; Censor, Horovitz, & Cohen, 2014; Dayan, Laor-Maayany, & Censor, 2016; Wymbs et al., 2016). It has been reported that existing motor memories can be modified through reconsolidation, which may repeat as needed across the lifespan (Censor, Horovitz, et al., 2014; Sandrini, Censor, Mishoe, & Cohen, 2013; Wymbs et al., 2016).

Measuring motor skill learning is not a trivial task. Most motor skills require the optimization of a speed-accuracy trade-off dependent upon specific task constraints. One approach to estimating learning is to reduce this feature to a single dimension by instructing participants to favor one factor over the other, or employing strict accuracy- or speed-related task requirements. An alternative approach is to use more neutral instructions or employ tasks that allow for natural variation of this interaction across the study population. In this case, the speed-accuracy trade-off is then explicitly modeled in performance or skill learning estimates. Another crucial factor in the experimental study of motor learning is the information participants have access to about the task and their performance. The specific nature and resolution of information available to participants will determine if learning is driven by factors such as sensory feedback error signals, cognitive strategies, or reward maximization. Thus, variants of the same basic task may assess very different learning processes. This may be particularly important for a technique like tDCS (which may exert its effect through the alteration of thresholds for neuronal discharge (Fritsch et al., 2010)) as observed effects may be highly dependent on the specific context in which it is applied.

Currently, the most frequently used tasks to investigate motor skill learning in experimental settings are: (1) sequential finger tapping tasks (SFTT; which can include either implicit or explicit sequence structure) (Ghilardi, Moisello, Silvestri, Ghez, & Krakauer, 2009; Nitsche et al., 2010; Reis et al., 2015; Song & Cohen, 2014); and (2) the sequential visual isometric pinch force task (SVIPT) (Reis et al., 2009). In a sense, these tasks are complimentary in that for the SFTT, the main unit of action is rather trivial for a healthy subject to accomplish (i.e. – pressing a keyboard key or button), while the required sequence of actions are typically complex in structure (between 8-15 items in length with controls on smaller intra-sequence patterns). Alternatively, the SVIPT requires execution of a precision pinch force action that is more difficult to elicit than a key-press (Waters-Metenier, Husain, Wiestler, & Diedrichsen, 2014). Thus, there is a greater emphasis placed upon accurate performance of the unit action within an explicit sequence context in the SVIPT than most variants of SFTTs. A general advantage of these learning tasks is that their complexity can be manipulated in a manner conducive to studying learning over long time periods (i.e. – months and years). Furthermore, competing sequences can be used to investigate consolidation and re-consolidation processes, as well.

Adaptation of highly-learned, target-directed pointing or shooting movements to environment perturbations has been regularly investigated (Orban de Xivry & Shadmehr, 2014). In this case, visual or proprioceptive feedback of generated movements is manipulated to produce a large error between a motor plan and sensory feedback. This error signal elicits an adaptive response that returns performance to pre-perturbation levels (Shadmehr, Smith, & Krakauer, 2010). The applied perturbations can be designed to affect either limb kinematics or dynamics, and typically involve rotating visual feedback representations of the movement (Krakauer, Ghilardi, & Ghez, 1999) or applying external forces to the moving limb via a robotic manipulandum (Smith, Brandt, & Shadmehr, 2000), respectively. As these tasks involve basic reaching movements that have been highly learned over a participant’s lifetime, performance levels typically return to baseline within a single training session.

Motor skill learning and adaptation are associated with functional and structural changes to a distributed brain network that includes primary motor (M1) and somatosensory (S1), dorsal (PMd) and ventral premotor (PMv), supplementary motor (SMA) and posterior parietal cortex (PPC), as well as the cerebellum and basal ganglia (Landi, Baguear, & Della-Maggiore, 2011; Pascual-Leone, Grafman, & Hallett, 1994; Scholz, Klein, Behrens, & Johansen-Berg, 2009). Thus, several candidate brain networks are accessible to tDCS or rTMS for investigating neuromodulatory effects on different features of motor learning. Furthermore, NIBS techniques are crucial for demonstrating that specific networks play an antecedent role in learning, as opposed to functional changes that emerge as a consequence (Diedrichsen & Kornysheva, 2015). To date, the primary region of interest for modulating online learning and retention of skill acquisition has been the contralateral, ipsilateral or bilateral M1 (Figure 2). In some cases, montages with electrodes positioned over PMd or the cerebellum have also been used, with cerebellum montages primarily used in relation to adaptation learning (Table 1). While tDCS electrodes have been placed overlying specific scalp locations, it should not be assumed that the underlying brain region is independently, specifically or selectively stimulated (Woods et al., 2016). Additionally, tDCS can modulate different stages of learning, best tested over multiple days.

**Figure 2.**
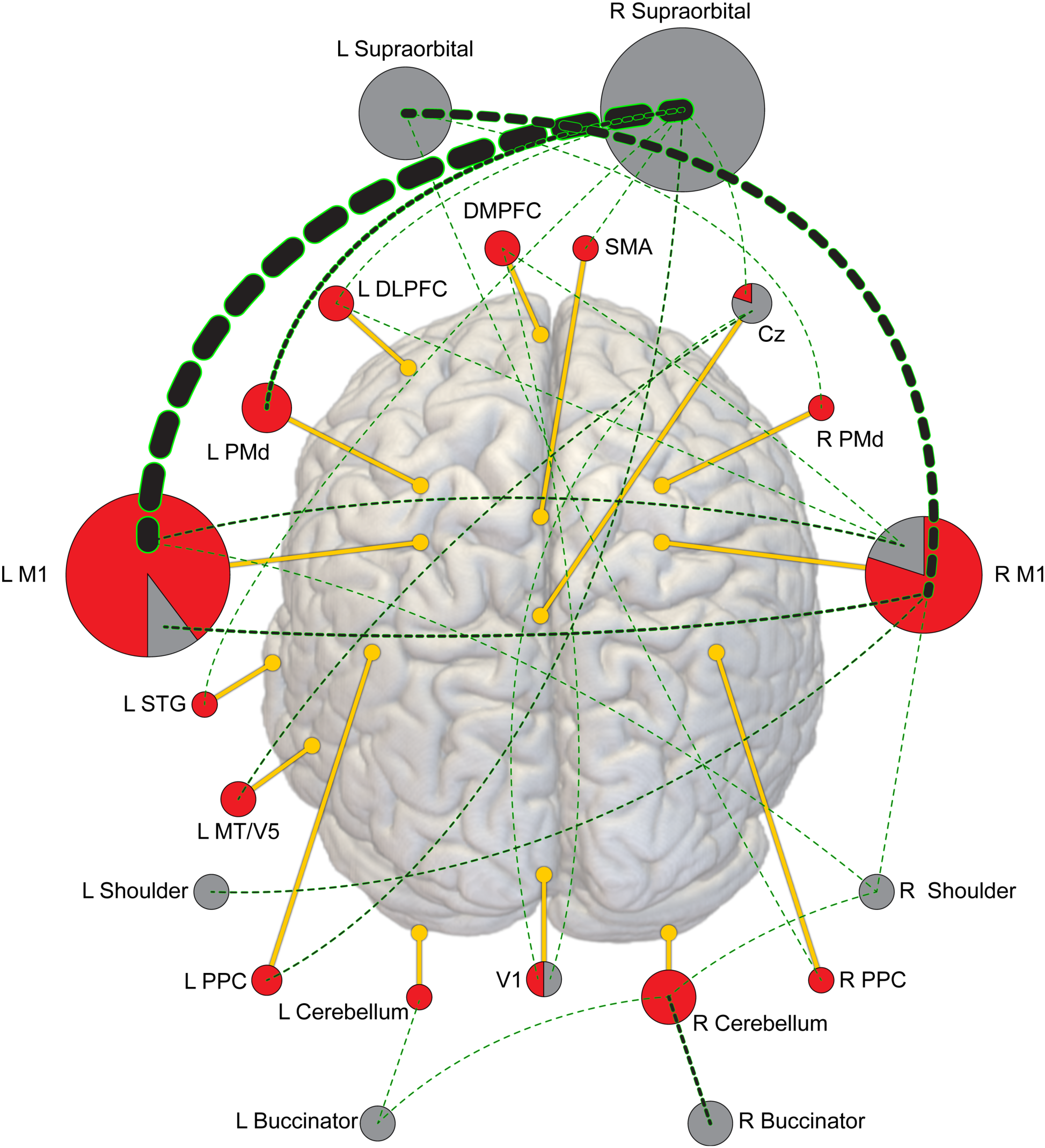
**Proportion of** tDCS montages utilized across eighty-three motor learning experiments. Circles for each location are proportionally filled with red (anode) and grey (cathode) to represent the relative number of studies the anode or cathode was placed at that location (i.e. –filled red circles were used as anode locations only, grey circles as cathode locations only, red and grey pie charts as both). The diameter of each circle is relative to the proportion of experiments that location was used in. Dashed lines represent montage connections between anode and cathode, with the line weighted relative to the proportion of experiments that particular montage was used in.

**Table 1.**
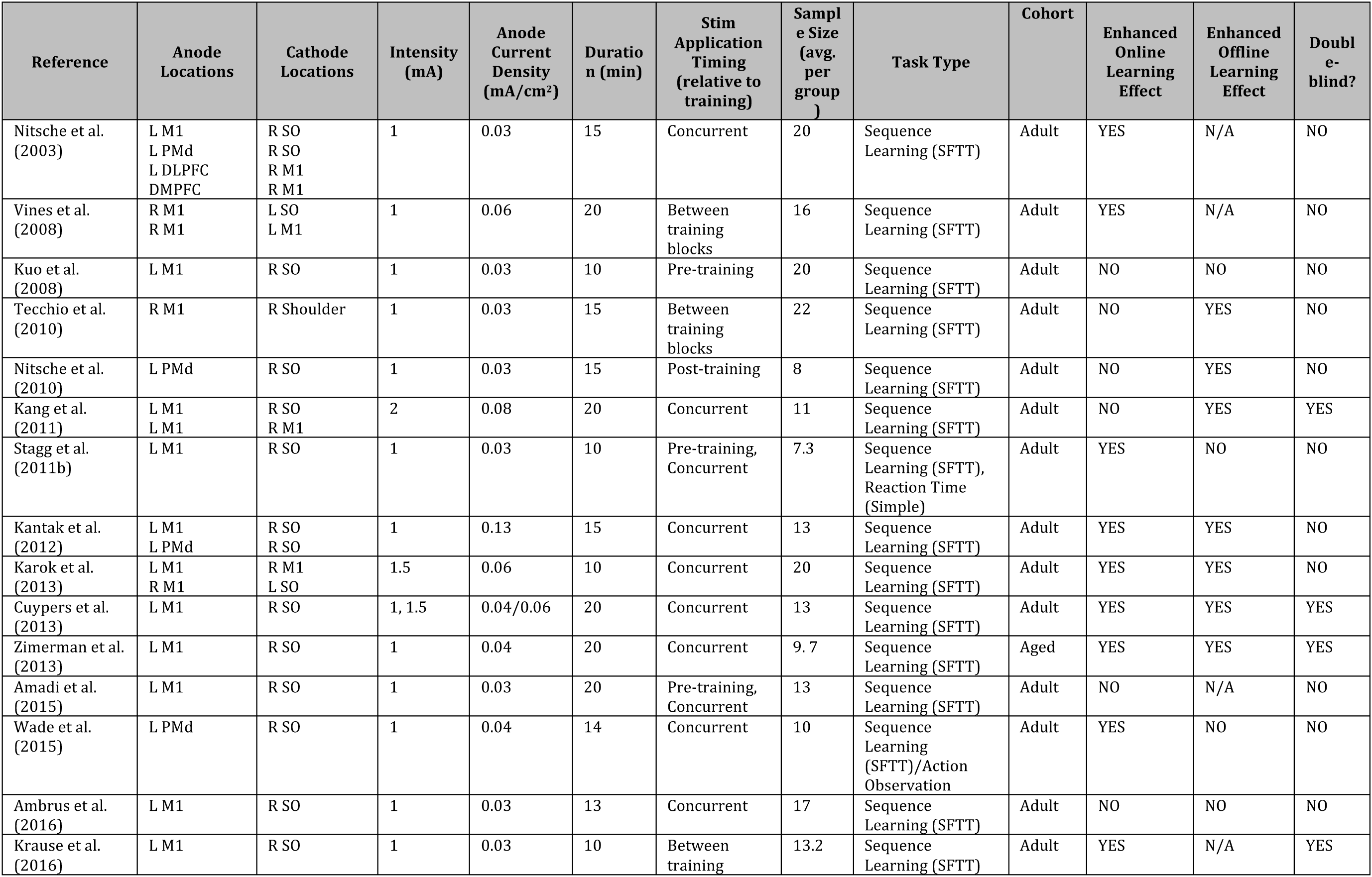
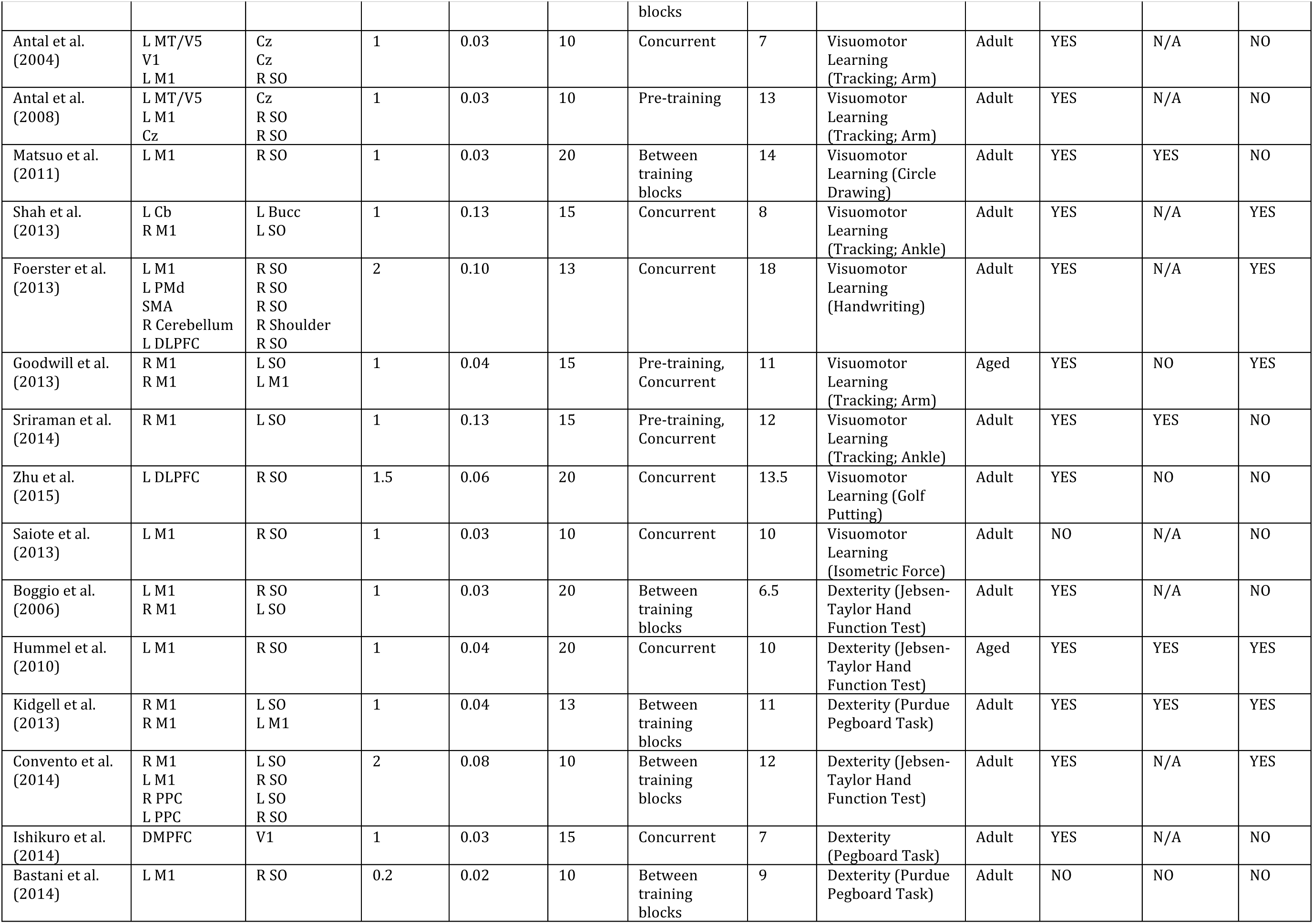
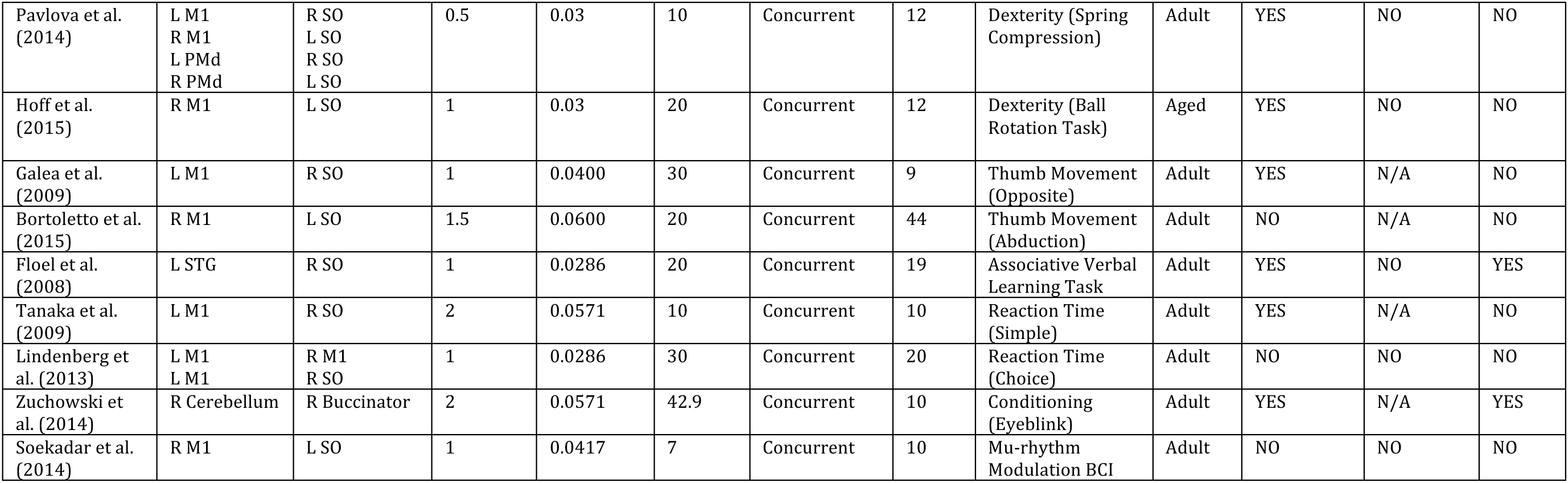
Investigations of tDCS-based enhancement of motor skill acquisition over a single day of training

### 1.2 Online motor performance and skill learning

Investigation of tDCS effects on online motor skill learning, that is performance gains observed during training, has focused primarily on stimulation montages where the anode is applied to a region of the scalp overlying the M1 contralateral to the practice hand and the cathode placed on the supraorbital (SO; sometimes referred to as “unilateral”) or M1 (sometimes referred to as “bilateral”) region of the opposite hemisphere. Other montages used have included anode placement over the cerebellum, PMd, PPC (specifically area MT/V5), and dorsomedial (DMPFC) or dorsolateral prefrontal cortex (DLPFC) (for more complex tasks) (Table 1). Stimulation parameters across these studies included median intensities of 1mA (range = 0.2– 2mA), target current densities of 0.04 mA/cm^2^ (0.0167–0.1327mA/cm^2^) and durations of 15 minutes (7–42.9 minutes). tDCS montage information and stimulation parameters for individual studies can be found in Tables 1-3. More remains to be learned about optimal parameters for eliciting specific behavioral effects. For example, it has been reported that tDCS applied with the same polarity may have opposing motor cortical excitability effects for different stimulation intensities (Batsikadze, Moliadze, Paulus, Kuo, & Nitsche, 2013). However, as different behavioral tasks have been employed in these studies the relationship between neurophysiological changes and resulting behavioral changes (which may vary across task domains) remains uncertain (López-Alonso, Cheeran, & Fernández-Del-Olmo, 2015).

The effects of tDCS on online sequence learning have been a particular area of interest (Amadi, Allman, Johansen-Berg, & Stagg, 2015; Ambrus et al., 2016; Cuypers et al., 2013; Kang & Paik, 2011; Kantak, Mummidisetty, & Stinear, 2012; Karok & Witney, 2013; M. F. Kuo et al., 2008; Nitsche et al., 2010; Nitsche et al., 2003; Reis et al., 2015; Reis et al., 2009; Stagg, Jayaram, et al., 2011; Tecchio et al., 2010; Vines, Cerruti, & Schlaug, 2008; Wade & Hammond, 2015). In an initial study, Nitsche and colleagues (2003) showed that tDCS with the anode applied over M1 concurrently with training improved online implicit learning of a motor sequence, while stimulation applied to PMd, DMPFC and DLPFC did not (Nitsche et al., 2003). Similar effects have been reported for explicit sequence learning, which may be GABA-mediated (Amadi et al., 2015; Stagg, Bachtiar, et al., 2011; Stagg, Jayaram, et al., 2011). Interestingly, stimulation with the anode applied over M1 prior to training appears to decrease subsequent learning rates (Amadi et al., 2015; Stagg, Bachtiar, et al., 2011; Stagg, Jayaram, et al., 2011), although whether or not this is mediated through a meta-plastic or homeostatic effect remains unclear (M. F. Kuo et al., 2008). Kantak and colleagues (2012) attempted to further dissociate tDCS-related effects for explicit versus implicit learning (Kantak et al., 2012). In this study, tDCS was delivered with the anode applied over M1 or PMd based upon previous work regarding the relative roles these areas play in explicit (where PMd is highly critical) versus implicit (where M1 is highly critical) learning. Stimulation delivered with the anode applied over M1 during an implicit motor sequence task resulted in greater online improvements compared with sham, as well as greater retention 24 hours later. In contrast, tDCS applied with the anode over PMd showed no online effects relative to sham, but in fact impaired retention at 24 hours. Finally, Cantarero et al. (2015) showed that applying tDCS stimulation with the anode over the ipsilateral cerebellum during skill learning (SVIPT task) in young healthy individuals augmented online skill acquisition via a reduction in error rates (Cantarero et al., 2015). This effect appeared to be robust, as it was present in every session for three consecutive days (the duration of the study). Interestingly, there were larger offline declines in the group receiving this type of tDCS, possibly due to a reduction in memory stability or that there was more accumulated knowledge to be lost. Despite this, the overall skill gains remained larger at one week follow up (Cantarero et al., 2015). Other previous work has shown significant online learning enhancement effects of tDCS applied with the anode located over M1 for early training sessions only (Reis et al., 2009).

Online tDCS-mediated effects for visuomotor skill learning (non sequential) for both the upper (Antal, Begemeier, Nitsche, & Paulus, 2008; Antal et al., 2004; Foerster et al., 2013; Matsuo et al., 2011; Zhu et al., 2015) and lower (Shah, Nguyen, & Madhavan, 2013; Sriraman, Oishi, & Madhavan, 2014) limb have also been investigated. Earlier work by Antal and colleagues (2004), showed that stimulation with the anode located over contralateral M1 or area MT/V5 (an extrastriate area that has been implicated in motion processing) applied during learning improved performance in a visuomotor tracking task when applied concurrently with training (Antal et al., 2004). Application of tDCS on these locations is consistent with known parietofrontal networks involved in these behaviors (Johnen et al., 2015). Using a naturalistic golf-putting task, Zhu et al. (2015) observed that cathodal stimulation applied over left DLPFC indirectly improved putting performance relative to sham stimulation (Zhu et al., 2015). This effect was particularly pronounced when participants were subjected to a multi-tasking constraint where putting and verbal working memory tasks were performed simultaneously. Overall, these results suggest that secondary network effects of stimulation (i.e. – alteration of information processing within the set of interconnected cortical areas) may play a more significant role in real-world environments where different cognitive and learning processes constantly interact.

Online training-induced improvements in non-dominant hand dexterity on the Purdue Pegboard and Jebsen-Taylor tests can be facilitated by tDCS (Bastani & Jaberzadeh, 2014; Convento, Bolognini, Fusaro, Lollo, & Vallar, 2014; Kidgell, Goodwill, Frazer, & Daly, 2013). Kidgell and colleagues (2013) found that tDCS applied with the anode over the non-dominant M1 using a unilateral (cathode over contralateral SO) or bilateral (cathode over contralateral M1) montage resulted in similar dexterity improvements (assessed with the Purdue Pegboard Test) in the non-dominant hand compared with sham stimulation (Kidgell et al., 2013). Using the same task, Bastani & Jaberzadeh (2014) investigated the effect of repeated offline application (up to 3) of relatively low intensity (0.2 mA) and duration (10 min) tDCS with the anode applied to the dominant (left) M1 on corticospinal excitability and behavior (Bastani & Jaberzadeh, 2014). Not surprisingly, given our understanding of the need for synchronous application of tDCS with training (Fritsch et al., 2010), no behavioral effects were observed. Of note however, corticospinal excitability was significantly facilitated up to 24 hours depending on the interval between subsequent stimulation applications, which had been reported previously (Monte-Silva et al., 2013). This finding suggests that cumulative effects of stimulation may be sensitive to the time between tDCS application and training. It also underscores the lack of a clear relationship between the neurophysiological and the behavioral effects of tDCS—changes in one may not predict or reflect changes in the other. Convento et al. (2014) found that offline tDCS with the anode applied to contralateral non-dominant M1 or ipsilateral PPC resulted in improved dexterity function in the non-dominant hand as well, with PPC and M1 stimulation having specific effects on action planning and execution, respectively (Convento et al., 2014).

A series of studies have looked at online learning and adaptation effects over the life span (Goodwill, Reynolds, Daly, & Kidgell, 2013; Hardwick & Celnik, 2014; Hoff et al., 2015; Hummel et al., 2010; Zimerman et al., 2013). Hummel and colleagues (2010) investigated motor performance effects of tDCS using a crossover design in a cohort of older adults (Hummel et al., 2010). When real or sham tDCS with the anode over the contralateral M1 was applied concurrently with performance of the Jebsen-Taylor hand function test (JTT) they observed a significant performance improvement relative to sham that lasted for over 30 min, and that the size of the effect correlated positively with age. The final group performance levels of the cohort were similar to those observed previously in a group of healthy young subjects. A later study by Zimerman and colleagues (2013) using very similar stimulation parameters looked instead at the effects of tDCS with the anode applied over M1 on sequence learning in aged adults (Zimerman et al., 2013). Again, performance gains were observed when the stimulation was applied to contralateral M1 concurrently with training, with effects remaining significant up to 24 hours later. More recently, it was reported that tDCS can influence learning in children (Ciechanski & Kirton, 2016)

Goodwill and colleagues (2013) assessed whether there was a differential effect of contralateral versus bilateral M1 stimulation with concurrent training on an upper limb visuomotor tracking task (Goodwill et al., 2013). Here, the group of older adults displayed similar performance gains and increased learning rates were observed for both contralateral and bilateral M1 stimulation, relative to sham. Furthermore, both montages resulted in the facilitation of corticospinal excitability and a decrease in observed short-interval intracortical inhibition (SICI). Complementary work by Zimerman and colleagues (2014) reported that cathodal stimulation applied to M1 ipsilateral to the learning hand actually impaired learning (Zimerman, Heise, Gerloff, Cohen, & Hummel, 2014). Finally, a recent study by Hardwick and Celnik (2014) compared the effects of tDCS with the anode located over the ipsilateral cerebellum between healthy younger and older individuals during a visuomotor adaptation (screen cursor rotation) task (Hardwick & Celnik, 2014). As expected, the group of older adults showed slower adaptation rates compared to younger adults when receiving sham tDCS. Older participants who received real tDCS, however, displayed faster learning rates that were similar in magnitude to the young group.

In summary, these studies suggest that tDCS applied with the anode over M1 or the cerebellum in a single training session may have broad-ranging effects on sequence learning or skill learning, respectively. Furthermore, this stimulation may be an effective tool in facilitating motor learning and adaptation in older healthy adult populations. It should be noted however, that in some studies online improvements are not seen, such as in several of the studies conducted over multiple days that emphasize offline effects (see Table 2).

**Table 2.**
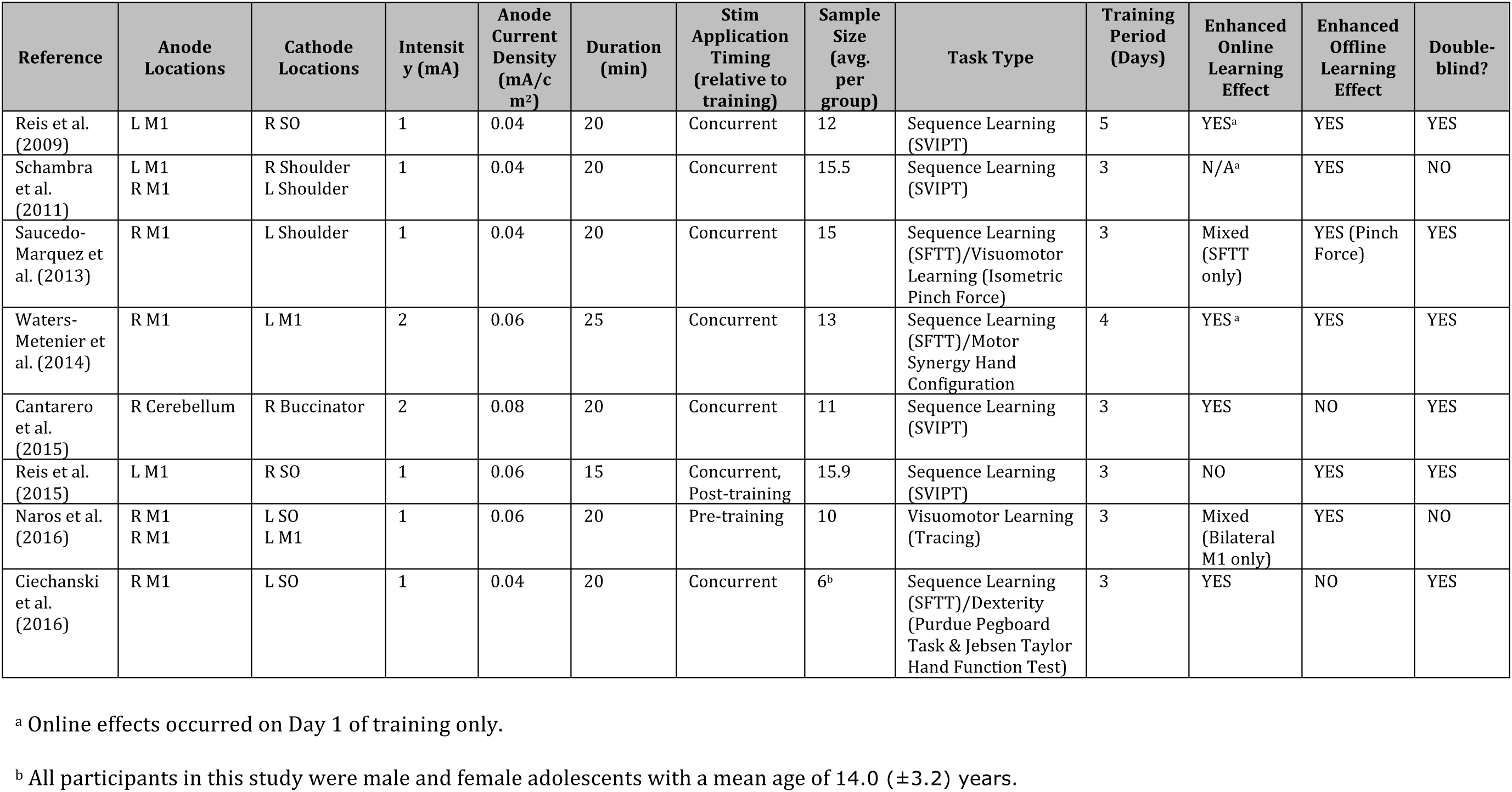
Studies investigating tDCS-based enhancement of motor skill learning and retention over multiple days of training

### 1.3 Offline motor skill learning and retention

Several studies, a majority of which have focused on sequence learning, have investigated offline motor skill learning and retention over multiple (typically at least three) days of training (Table 2) (Cantarero et al., 2015; Naros et al., 2016; Reis et al., 2015; Reis et al., 2009; Saucedo-Marquez, Zhang, Swinnen, Meesen, & Wenderoth, 2013; Schambra et al., 2011; Waters-Metenier et al., 2014). Reis and colleagues (2009) found that tDCS with the anode located over M1 applied concurrently with training over five consecutive days resulted in significant enhancement of offline skill gains and retention compared with sham in the sequential visual isometric pinch-force task (SVIPT) (Reis et al., 2009). While learning within sessions was not significantly different between the two groups, learning over the five sessions was facilitated in the group receiving real stimulation. Furthermore, this difference remained present when skill was retested three months later, suggesting that these gains had successfully consolidated and remained stable over the long-term. In a follow-up study, effects mediated by consolidation processes were further supported, as offline skill gains induced by real tDCS were found to be dependent upon the passage of time as opposed to requiring overnight sleep (Reis et al., 2015). Concurrent application of tDCS with training also appears crucial for these effects to emerge as stimulation applied post-training only did not induce offline skill gains, consistent with the finding that tDCS alone does not elicit LTP unless it is associated with a second input delivered to the motor cortex in rodents (Fritsch et al., 2010). Modifications made to the montage used here (cathode placed over contralateral supraorbital location) to an alternative montage with extracephalic cathode location (ipsilateral shoulder) resulted in reduced effects of stimulation (Schambra et al., 2011). In agreement with modeling predictions, this finding suggests that the montage configuration is the primary determinant of the applied current density distribution, and plays an important role in resulting behavioral effects (Bestmann, 2015; de Berker, Bikson, & Bestmann, 2013; Woods et al., 2016). Finally, tDCS applied with the anode located over the cerebellum increased skill learning in this task through the enhancement of online as opposed to offline components. In particular, the larger gains were driven to a greater extent by reductions in error rates as opposed to changes in movement time. This suggests that specific task constraints may play a role in determining the motor network areas of interest (Cantarero et al., 2015). For example, tDCS with the anode placed over the cerebellum applied concurrently with training for a task with very precise timing requirements enhanced offline improvement, as opposed to online learning as observed in prior studies (Wessel et al., 2016).

Saucedo-Marquez and colleagues (2013) conducted a crossover design that investigated the decomposable elements of the SVIPT task, sequence learning (sequential finger tapping) and visual isometric pinch force (Saucedo-Marquez et al., 2013). Following three days of training in each task with concurrent application of sham or real tDCS applied with the anode over M1, they observed that real stimulation improved online sequence learning, but only skill retention for the pinch force task. In addition to task-specific learning effects, these findings suggest that different learning processes interact with tDCS stimulation in non-additive ways as task complexity increases. Waters-Metenier et al. (2014) looked at task-specific effects of bilateral M1 stimulation, in this case on the learning novel hand movement synergy patterns and finger-tapping sequences (Waters-Metenier et al., 2014). In this case, tDCS improved both synergy and sequence learning with long-term retention of the effects persisting for at least 4 weeks following training. Furthermore, bilateral M1 stimulation effects showed task- and effector-based generalization to untrained hand synergies and finger sequences, and the untrained hand, respectively. This generalization is most likely the result of polarity specific effects on each hemisphere (Naros et al., 2016).

### 1.4 Adaptation

tDCS-related effects on adaptation have also been studied in young healthy adults (Avila et al., 2015; Galea et al., 2011; Herzfeld et al., 2014; Hunter, Sacco, Nitsche, & Turner, 2009; Orban de Xivry et al., 2011) (Table 3). Galea and colleagues (2011) compared the effects of tDCS applied with the anode over the cerebellum versus M1 during concurrent adaptation to 30-degree rotation of visual feedback (Galea et al., 2011). Here, cerebellar tDCS resulted in faster initial adaptation to the perturbed task environment, while M1 stimulation showed no effect in this regard. In contrast, a dissociative effect emerged when M1 stimulation resulted in improved retention of the newly acquired visuomotor transformation, as subjects receiving this stimulation adapted faster when the perturbation was reintroduced following a washout period. Interestingly, in a force-field reaching task that assesses adaptation to perturbed upper limb dynamics, tDCS applied with the anode over the cerebellum increased error-dependent learning and facilitated adaptation, while M1 stimulation had no effect (Herzfeld et al., 2014). Furthermore, tDCS applied with the anode over M1 did not improve retention. In addition to the work above, this suggests that M1 and the cerebellum play complementary roles with respect to different learning processes, and that tDCS can be used to influence these processes in a task-dependent manner.

**Table 3.** Studies investigating tDCS-based enhancement of motor skill adaptation in healthy subjects

| Reference | Anode Locations | Cathode Locations | Intensity (mA) | Anode Current Density (mA/cm<sup>2</sup>) | Duration (min) | Stim Application Timing (relative to training) | Sample Size (avg. per group) | Task Type | Cohort | Enhanced Online Learning Effect | Enhanced Offline Learning Effect | Double-blind? |
| --- | --- | --- | --- | --- | --- | --- | --- | --- | --- | --- | --- | --- |
| Hunter et al. (2009) | L M1 | R SO | 1 | 0.03 | 17 | Concurrent | 14 | Adaptation (Force-field) | Adult | NO | NO | NO |
| de Xivry et al. (2011) | L M1<br>L PPC | R SO<br>R SO | 1 | 0.04 | 20 | Concurrent | 8.6 | Adaptation (Force-field) | Adult | YES | NO | NO |
| Galea et al. (2011) | L M1<br>R Cerebellum | R SO<br>R Buccinator | 2 | 0.08 | 15 | Concurrent | 9.5 | Adaptation (Screen Cursor Rotation) | Adult | YES (Cerebellum) | YES (M1) | YES |
| Herzfeld et al. (2014) | L M1<br>R Cerebellum | R SO<br>R Buccinator | 2 | 0.08 | 25 | Concurrent | 12.75 | Adaptation (Force-field) | Adult | YES | NO | YES |
| Hardwick & Celnik (2014) | R Cerebellum | R Buccinator | 2 | 0.08 | 15 | Concurrent | 11 | Adaptation (Screen Cursor Rotation) | Aged | YES | NO | NO |
| Avila et al. (2015) | R Cerebellum | L Buccinator | 1.5 | 0.13 | 15 | Pre-training, Concurrent | 10 | Adaptation (Saccade) | Adult | YES | NO | YES |

### 2 Meta-analyses and systematic reviews of the literature

Hashemirad and colleagues (Hashemirad et al., 2016) reviewed the effects of tDCS with the anode placed over M1 on motor sequence learning in healthy adults. 13/140 reviewed articles (9.2%) met the eligibility criteria (one or more sessions of unilateral or bilateral tDCS over M1 concurrently with training the SFTT or SVIPT tasks, and included a negative control group for stimulation (either sham tDCS plus task training or training only)). The authors conclude that the effects of tDCS applied with the anode over M1 on sequential motor learning may depend on learning stages (Dayan & Cohen, 2011) and be to some extent task- or montage-specific (Schambra et al., 2011) and that multiple tDCS sessions present advantages over single session applications on both finger tapping and SVIPT tasks. Similarly, the effects on long-term retention might be task specific with different retention effects reported in the finger tapping versus SVIPT tasks (Reis et al., 2015; Reis et al., 2009; Saucedo-Marquez et al., 2013; Waters-Metenier et al., 2014). Of note, the relatively small number of studies fitting the inclusion criteria is a clear illustration of the challenges faced when attempting to perform quantitative reviews of tDCS effects on motor learning (Antal, Keeser, Priori, Padberg, & Nitsche, 2015; Nitsche, Bikson, & Bestmann, 2015). Other meta-analyses focusing on effects of a single tDCS session have reported few significant physiological (Horvath, Forte, & Carter, 2015a) and no significant cognitive effects (Horvath, Forte, & Carter, 2015b), although questions regarding methodology used in these analyses have been raised (Antal et al., 2015). Additionally, it should be kept in mind that in the absence of systematic critical assessment of the quality of individual studies, and understanding of the biases that they may be prone to, interpretation of meta-analysis findings remains uncertain (Bastian, 2016).

## 3 Caveats and considerations for the future

There has been a substantial increase in the number of investigations using tDCS over M1 to influence motor learning since the previous consensus document in 2008 (Reis et al., 2008). Since then, a number of scientific, methodological and social caveats have emerged that deserve closer scrutiny by those interested in using this technique. Many of these caveats are applicable to the broader realm of basic and clinical science, while others are more specific to the use of tDCS.

Scientific caveats include understanding that application of tDCS with one of the electrodes placed over a specific region may not influence that region or may result in behavioral changes through distant (i.e. - poor spatial targeting or focality) or secondary effects on other interconnected cortical areas (Dayan et al., 2013), infrequent use of modeling to guide stimulation montages (de Berker et al., 2016) or overly simplified modeling assumptions that neglect the folding of the cortex and consequences on stimulation effects (i.e. – decreasing the threshold for hyperpolarization of neurons on one side of a gyrus but depolarization on the other). Systematic determination of the optimal timing of stimulation for inducing long-lasting effects, and how this varies across individuals, is another avenue where more research is needed (Manenti, Sandrini, Brambilla, & Cotelli, 2016; Martin, Liu, Alonzo, Green, & Loo, 2014). Indeed, a more coordinated effort where experimental parameters and modeling assumptions are iteratively refined is required (Bestmann, 2015; Brunoni et al., 2012).

While the neuromodulatory after-effects induced by NIBS techniques (including tDCS) appear to be relatively stable over prolonged time courses (López-Alonso et al., 2015), the nature and magnitude of these effects varies considerably between individuals (Hamada, Murase, Hasan, Balaratnam, & Rothwell, 2013; Nettekoven et al., 2015; Nitsche & Paulus, 2001; Wiethoff, Hamada, & Rothwell, 2014). One source of this variability may be the brain state-dependent nature of these effects, meaning that the history of endogenous activity of one region may be crucial to the effects of brain stimulation (Silvanto et al., 2008) and consequent activation of homeostatic and non-homeostatic metaplasticity mechanisms (Amadi et al., 2015; Muller-Dahlhaus & Ziemann, 2015). More complete investigation of these proposed factors represents an important hurdle for elucidating inter-individual variability. Furthermore, under-reporting of negative effects (Horvath et al., 2015b) due to publication bias (Mancuso, Ilieva, Hamilton, & Farah, 2016; Shiozawa et al., 2014; Vannorsdall et al., 2016) represents another important scientific caveat that must be addressed in order to facilitate future research progress.

Motor learning is a rather complex process in itself, with different forms (i.e.- use-dependent, error-based, reinforcement, strategic learning) and likely different underlying neural substrates. Many of the tasks employed to determine the effects of tDCS on learning either have several variants or include different forms of learning. These circumstances limit the information that can be drawn from the effects of tDCS on those tasks. For instance, it is possible that tDCS changes learning because it improves knowledge of the dynamics of the task at hand, or because it improves the strategic approach to that task. Depending on the specific task variant or learning strategies employed by a given individual, tDCS applied to one region may or may not influence task learning. Therefore, a better understanding of motor learning processes, and the tasks used to assess them, will be critical to determine whether NIBS can or cannot manipulate behaviors that are potentially impactful to daily life. Similarly, the issue of generalization is of clear relevance to rehabilitation and remains a major challenge. In addition to investigating the efficacy of tDCS in enhancing specific quantitative features of skill learning, improving our understanding of the effects of tDCS on the generalization of learning across different skills will also be an important scientific endeavor (Waters-Metenier et al., 2014).

Maturation of the tDCS field since the previous consensus document (Reis et al., 2008) and the focus on enhancing human motor learning have overall raised the bar of methodological and design requirements in tDCS studies. Present problems in the field include: (1) insufficient use of double-blind designs (see above, for example only 25 out of the 60 published studies on tDCS effects on motor learning in healthy adults reviewed here utilized double-blind designs) and positive controls (stimulation of other cortical regions); (2) insufficient differentiation and understanding of design and claims when carrying out exploratory (hypothesis-generating) versus confirmatory (hypothesis-driven) research (the former suggesting trends and providing data for prospective power analysis and the latter, strengthened by preregistration (Finkel, Eastwick, & Reis, 2015), allowing drawing conclusions on particular effects; (3) insufficient efforts to reduce false-positive rates in studies geared to provide proof of principle data to power subsequent clinical trials; (4) scarcity of preregistration of hypothesis, design, power analysis and data processing for research written up as hypothesis-driven and confirmatory (see for example https://blogs.royalsociety.org/publishing/registered-reports/); (5) insufficient prepublication and sharing of materials (Lauer, Krumholz, & Topol, 2015; Morey et al., 2016), particularly in relation to negative results; (6) insufficient post-publication repositories of data (see for example (Campbell et al., 2002)) and in general (Nosek et al., 2015)) to allow additional analyses; (7) seldom use of experimental designs with replications built in (Anderson et al., 2016; Cohen et al., 1997; Gilbert, King, Pettigrew, & Wilson, 2016; Nosek et al., 2015); and (8) use of appropriate sample size based on prospective power analysis for studies claimed to be hypothesis-driven.

Another important discussion surrounding tDCS research is related to how reproducibility of reported effects should be evaluated. A special mention should be made to the expression of the general reproducibility problem in science (Collins & Tabak, 2014) to tDCS studies of motor learning. There are three levels of reproducibility: methods, results and inferential (Goodman, Fanelli, & Ioannidis, 2016). Methodological reproducibility requires “*provision of enough detail about study procedures and data so the same procedures could …be exactly repeated*”. More importantly, in order to evaluate methodological reproducibility, there should be “… *agreement about the level of detail needed in the description of the measurement process, …the degree of processing of the raw data …”* and the “*completeness of the analytic reporting*”. Such agreement does not exist at the present time in the tDCS field. Development of standards of consistency in methodological reporting would represent an important step forward. To start addressing this problem, we propose a checklist with reporting standards for tDCS studies (Table 4). Reproducibility of results refers to replicability once the tools for methodological replication are fully provided and agreed upon. Importantly, replicability is best tested for stochastic data using Bayesian paradigms of accumulating evidence more than binary criteria of successful or unsuccessful replication (Goodman et al., 2016). Clearly, “*statistical significance by itself tells very little about whether one study has “replicated” the results of another*”. Finally, inferential reproducibility refers to “*drawing of qualitatively similar conclusions from either an independent replication of a study or a reanalysis of the original study*”. Please, see Goodman and Ioannidis for a full discussion (Goodman et al., 2016).

**Table 4.**
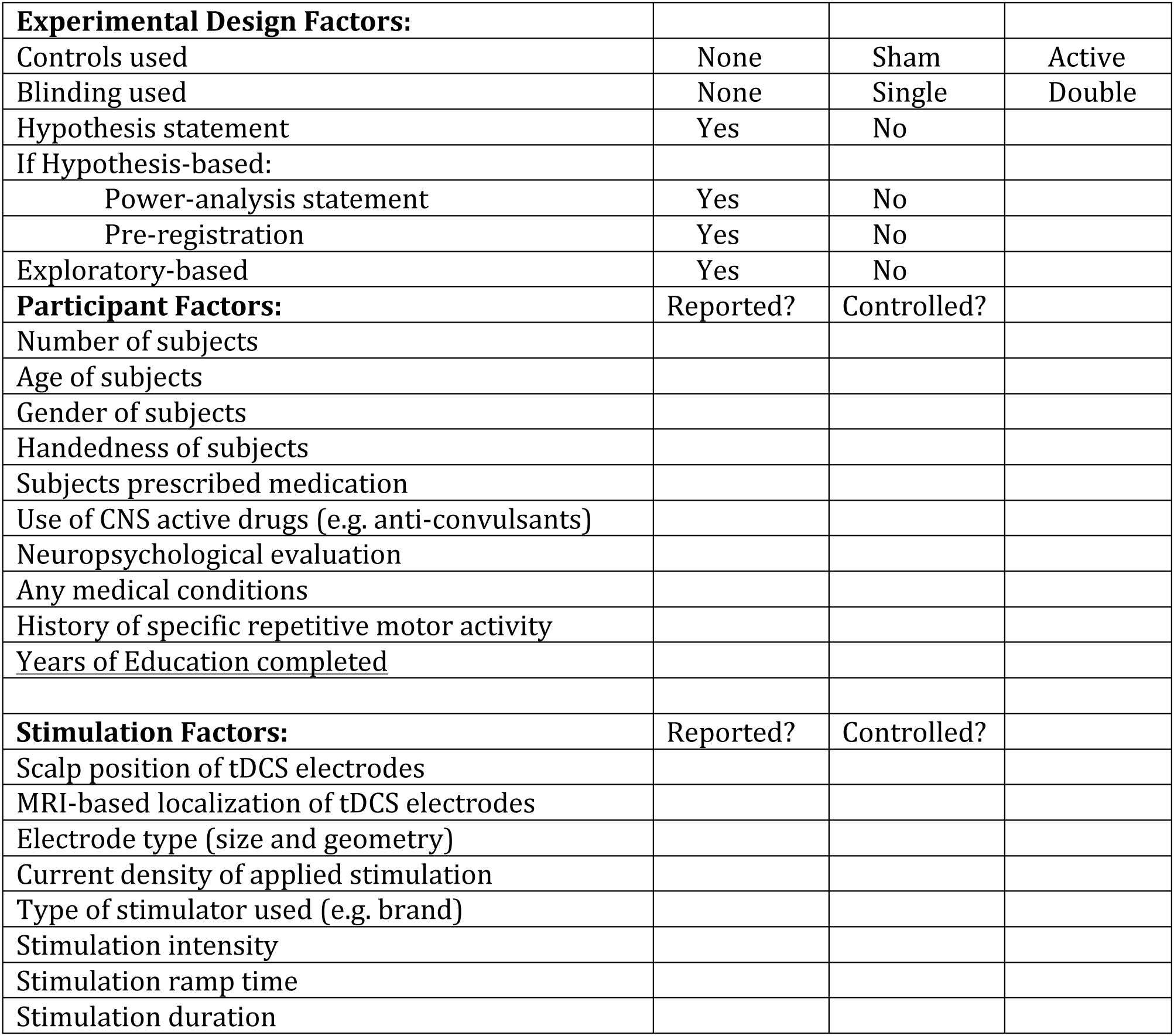
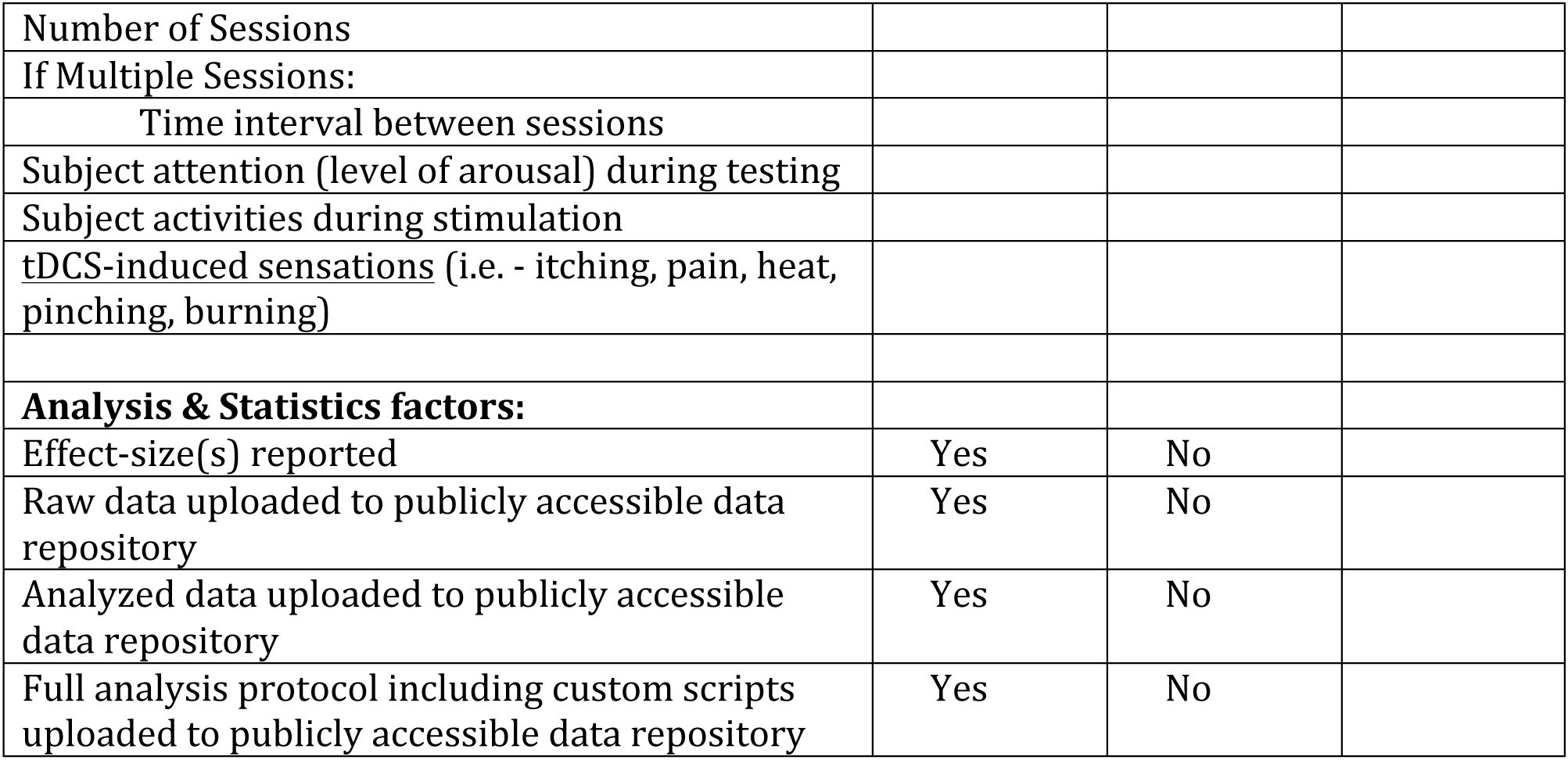
Reporting checklist for tDCS studies. Modified from (Chipchase et al., 2012)

|  |  |  |  |
| --- | --- | --- | --- |
| **Experimental Design Factors:** |  |  |  |
| Controls used | ☐ None | ☐ Sham | ☐ Active |
| Blinding used | ☐ None | ☐ Single | ☐ Double |
| Hypothesis statement | ☐ Yes | ☐ No |  |
| If Hypothesis-based: |  |  |  |
| Power-analysis statement | ☐ Yes | ☐ No |  |
| Pre-registration | ☐ Yes | ☐ No |  |
| Exploratory-based | ☐ Yes | ☐ No |  |
| **Participant Factors:** | Reported? | Controlled? |  |
| Number of subjects | ☐ | ☐ |  |
| Age of subjects | ☐ | ☐ |  |
| Gender of subjects | ☐ | ☐ |  |
| Handedness of subjects | ☐ | ☐ |  |
| Subjects prescribed medication | ☐ | ☐ |  |
| Use of CNS active drugs (e.g. anti-convulsants) | ☐ | ☐ |  |
| Neuropsychological evaluation | ☐ | ☐ |  |
| Any medical conditions | ☐ | ☐ |  |
| History of specific repetitive motor activity | ☐ | ☐ |  |
| Years of Education completed | ☐ | ☐ |  |
| **Stimulation Factors:** | Reported? | Controlled? |  |
| Scalp position of tDCS electrodes | ☐ | ☐ |  |
| MRI-based localization of tDCS electrodes | ☐ | ☐ |  |
| Electrode type (size and geometry) | ☐ | ☐ |  |
| Current density of applied stimulation | ☐ | ☐ |  |
| Type of stimulator used (e.g. brand) | ☐ | ☐ |  |
| Stimulation intensity | ☐ | ☐ |  |
| Stimulation ramp time | ☐ | ☐ |  |
| Stimulation duration | ☐ | ☐ |  |

|  |  |  |
| --- | --- | --- |
| Number of Sessions | ☐ | ☐ |
| If Multiple Sessions: |  |  |
| Time interval between sessions | ☐ | ☐ |
| Subject attention (level of arousal) during testing | ☐ | ☐ |
| Subject activities during stimulation | ☐ | ☐ |
| tDCS-induced sensations (i.e. - itching, pain, heat, pinching, burning) | ☐ | ☐ |
| **Analysis & Statistics factors:** |  |  |
| Effect-size(s) reported | ☐ Yes | ☐ No |
| Raw data uploaded to publicly accessible data repository | ☐ Yes | ☐ No |
| Analyzed data uploaded to publicly accessible data repository | ☐ Yes | ☐ No |
| Full analysis protocol including custom scripts uploaded to publicly accessible data repository | ☐ Yes | ☐ No |

A unique concern that has emerged with transcranial electrical stimulation techniques, is that the simplicity, low-cost nature of, and public access to the technology has lead to the emergence of a popular do-it-yourself movement where individuals participate in self-experimentation without oversight. Such data is not part of well-designed experimental protocols (Fitz & Reiner, 2015; Riggall et al., 2015; Wexler, 2016). An additional worrisome aspect of this movement is that no studies have investigated yet the long-term effects associated with tDCS chronic use (for further reading please see http://www.ifcn.info/uploadfiles/documents/2015/Using_tES_devices_as_DIY_FINAL_13Dec15.pdf).

As nuanced understanding of the possibilities and limitations of a given experimental technique matures, critical evaluation amongst experts leads to the progressive refinement of standards associated with its use. Used alone, tDCS has a quite large parameter space. On one hand, this flexibility is one of the main features supporting the general use of tDCS across several disciplines and purposes. However, this has resulted in substantial variation in stimulation parameters across individual studies and laboratories, and has presented a challenge to the convergence upon field-wide standards. Furthermore, when used in conjunction with different behavioral tasks (or even variants of a single task) this dimensionality substantially grows. An interesting approach to addressing the issue of heterogeneity of stimulation protocols and tasks could be to directly account for the heterogeneity within statistical models through inclusion of stimulation parameters, electrode montages and task variants as covariates. In this way, meta-analyses could serve as important tools for identifying which experimental factors predominantly explain significant levels of inter- or intra-individual variability(Horvath, Vogrin, Carter, Cook, & Forte, 2016; Lopez-Alonso, Fernandez-Del-Olmo, Costantini, Gonzalez-Henriquez, & Cheeran, 2015).

## 4 Conclusions

The 2008 consensus concluded: “*In summary, the scarce studies performed so far point to the encouraging conclusion that noninvasive brain stimulation can contribute to the understanding of mechanisms underlying motor learning and motor memory formation and raise the exciting hypothesis that this increased understanding could in the future result in the development of new strategies to enhance specific stages of learning and memory processing in healthy humans and in patients with brain lesions*”. A growing body of work continues to support the use of noninvasive brain stimulation as a tool for neuromodulation of motor learning. However, the larger literature has raised numerous and substantial caveats to be considered that are not trivial to resolve. More work is required to understand mechanisms underlying the effects of tDCS and substrates of inter-individual variability, to optimize dosing and methodological designs. Additionally, improved understanding of motor skill learning processes and standardization of tasks will help reduce inter-study variability, as the scientific approach to manipulating motor learning will become more precise. Emerging efforts for improving transparency, full reporting of data and all analyses carried out, replication and data sharing through repositories will be important to answering these questions.

## **Footnote** (p. 3; first paragraph of Introduction)

a TMS-based investigations have included the use of repetitive (primarily 1, 5 or 10Hz) and patterned (continuous or intermittent theta burst; cTBS or iTBS, respectively) stimulation protocols. Transcranial electrical stimulation (TES) techniques have included direct (tDCS) or alternating current (tACS) (Krause, Meier, Dinkelbach, & Pollok, 2016; Pollok, Boysen, & Krause, 2015), or random noise (tRNS) (Saiote, Polanía, Rosenberger, Paulus, & Antal, 2013) stimulation. Since published findings using rTMS, TBS, tACS and tRNS for enhancing motor learning remain particularly sparse (Figure 1) the primary focus of the review will be on tDCS-based interventions.

